# Dysregulation of the p16–CDK6 axis characterizes HPV-unrelated p16-positive oropharyngeal carcinoma

**DOI:** 10.64898/2026.08.17.743010

**Authors:** Kazuki Hayashi, Miki Kobayashi, Taku Kitano, Takahito Fukusumi, Toshihiro Kishikawa, Takashi Fujii, Rikifumi Ohta, Shinichi Morishita, Eiji Hara, Hidenori Inohara, Tomonori Matsumoto

## Abstract

Human papillomavirus (HPV)-related and HPV-unrelated oropharyngeal squamous cell carcinomas (OPCs) are distinct entities with different clinical outcomes. While p16 immunohistochemistry (IHC) is widely used as a surrogate marker for HPV-driven OPC, a subset of HPV-unrelated OPCs also overexpress p16, and the biological basis of this discordance remains unclear. Here, we performed integrated clinicopathological, transcriptomic, genomic, and functional analyses of OPCs and demonstrated that dysregulation of the p16–CDK6 axis characterizes HPV-unrelated p16-positive OPCs. Although these tumors closely resembled HPV-unrelated p16-negative OPCs in their clinicopathological and transcriptomic characteristics, they exhibited a more favorable prognosis. CDK6 was recurrently upregulated in HPV-unrelated OPC regardless of p16 status and was already detectable in high-grade dysplastic leukoplakia, suggesting that CDK6 activation is an early event in HPV-unrelated tumorigenesis. In experimental models, CDK6 overexpression induced compensatory p16 upregulation, creating selective pressure for subsequent CDKN2A inactivation. Consistent with this model, homozygous CDKN2A loss predominated in p16-negative tumors. We further identified CDKN2A frameshift mutations generating p14ARF–p16 chimeric proteins that retain p16 immunoreactivity despite functional loss of wild-type p16, revealing a previously unrecognized diagnostic pitfall of p16 IHC. These findings provide a biological framework for p16 overexpression in HPV-unrelated OPC and suggest that assessment of the p16–CDK6 axis may refine molecular classification and risk stratification beyond p16 IHC alone.

## Introduction

Oropharyngeal squamous cell carcinoma (OPC) is a heterogeneous disease that can arise through distinct etiological mechanisms, including exposure to alcohol and tobacco and infection with human papillomavirus (HPV) (1, 2). In recent decades, HPV-driven OPC has become increasingly prevalent, particularly in developed countries (3, 4). Importantly, HPV status has a profound impact on the clinical behavior of OPC. Patients with HPV-related OPC generally exhibit a more favorable prognosis than those with HPV-unrelated disease (3, 5, 6). Accordingly, assessment of HPV status is recommended for all newly diagnosed OPC, and contemporary staging systems explicitly distinguish HPV-related from HPV-unrelated OPC (7, 8). Treatment strategies are also being investigated in the context of HPV status, with de-escalation approaches aimed at reducing long-term toxicities actively explored for HPV-related OPC, while conventional treatment approaches remain the standard for HPV-unrelated OPC (6).

Although HPV-related and HPV-unrelated OPC arise through distinct molecular mechanisms, disruption of both the p53 and retinoblastoma (Rb) tumor suppressor pathways is a shared hallmark of carcinogenesis. In HPV-related tumors, the viral oncoproteins E6 and E7 functionally inactivate these pathways by promoting p53 degradation and Rb inactivation, respectively (3, 9, 10). In contrast, HPV-unrelated cancers typically harbor TP53 mutations together with mutation or deletion of CDKN2A, which encodes the cyclin-dependent kinase inhibitor p16, a key upstream regulator of the Rb pathway (11, 12). While disruption of both pathways is central to the development of OPC, alterations affecting the p16–Rb axis are particularly relevant to clinical classification because p16 overexpression is widely used as a surrogate marker of HPV-related disease.

In clinical practice, determination of HPV status in OPC is largely based on these mechanistic differences. Because E7-mediated inactivation of Rb induces p16 overexpression through a negative-feedback mechanism, p16 immunohistochemistry (IHC) is widely used as a practical surrogate for HPV association because of its simplicity and broad applicability (3, 4). However, accumulating evidence has shown that p16 IHC does not always concord with HPV detection by nucleic acid-based assays, including PCR and in situ hybridization for HPV DNA or mRNA (13–15). Approximately 10% of p16-positive OPCs are negative for HPV DNA or RNA (4, 13). Importantly, patients with p16+/HPV− OPC have outcomes intermediate between those with p16+/HPV+ and p16−/HPV− disease (13). These findings suggest the existence of a distinct subgroup of OPC characterized by p16 overexpression in the absence of HPV infection. However, whether p16 overexpression simply reflects discordance between p16 IHC and HPV status or instead represents a biologically meaningful state with implications for understanding HPV-unrelated OPC remains unresolved.

In this study, we sought to characterize the molecular features of HPV-unrelated p16-positive OPC and to determine the biological significance of p16 overexpression in this context. By integrating clinicopathological, transcriptomic, genomic, and functional analyses, we identify dysregulation of the p16–CDK6 axis as a characteristic feature of this subgroup. We further show that CDK6 upregulation emerges early during tumorigenesis and distinguishes HPV-unrelated from HPV-related OPC, supporting a model in which aberrant regulation of the p16–CDK6 axis contributes to HPV-unrelated OPC development.

## Results

### False-positive p16 immunostaining in HPV-unrelated cancers is associated with a better prognosis than p16 negativity

Among 352 patients with OPC identified during the enrollment period, 316 patients who underwent radical treatment and had evaluable p16 and HPV status were included in the analysis (Fig. 1A). In accordance with international guidelines (16, 17), all tumors were screened by p16 IHC using clinical specimens obtained at the initial visit (Fig. 1B). p16 immunostaining with the E6H4 clone was positive in 46.2% (146/316) of cases. By contrast, HPV PCR targeting the E6 and E7 regions revealed substantial discordance: 23 p16-positive cases were HPV PCR–negative. Overall, p16 immunostaining showed a sensitivity of 98.4% and a specificity of 88.0% for oncogenic HPV infection, with a positive predictive value of 84.2% (Fig. 1A). Thus, p16 IHC was highly sensitive for detecting oncogenic HPV infection but yielded false-positive results in a subset of cases.

**Figure 1.**
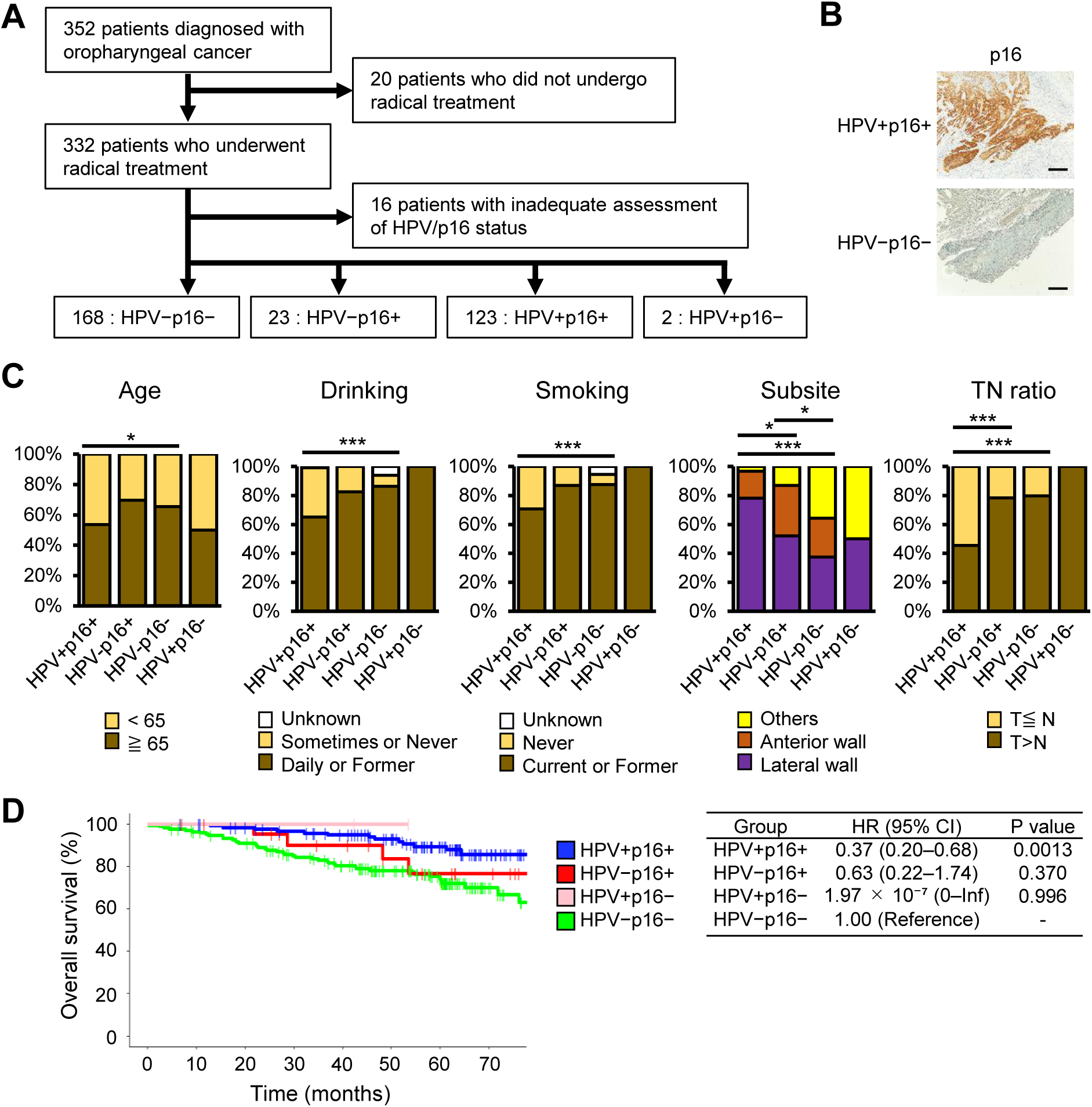
Clinical characteristics according to HPV/p16 Status in OPC. (A) Study flow diagram. Of the patients diagnosed with OPC at initial consultation, 20 who did not receive curative-intent treatment at the initial visit and 16 with unknown HPV/p16 status were excluded. (B) Representative images of p16 immunohistochemical staining using the E6H4 antibody. Scale bars, 200 μm. (C) Clinical characteristics of OPC according to HPV/p16 status. Only two cases were HPV+/p16– and were excluded from statistical analyses. Chi-square test. *p < 0.05, **p < 0.01, ***p < 0.001. (D) Kaplan–Meier estimates of overall survival stratified by HPV/p16 status. Hazard ratios (HRs) and P values were estimated using the Cox proportional hazards model. The HPV−/p16− group was used as the reference. CI, confidence interval; Inf, infinity.

We next compared the clinical characteristics of p16-positive/HPV-PCR–negative cases (hereafter “HPV-unrelated p16+”) with those of p16-positive/HPV-PCR–positive cases (“HPV-related”) and p16-negative/HPV-PCR–negative cases (“p16−”). Patients with p16− tumors were significantly older and had higher rates of alcohol consumption and smoking than those with HPV-related tumors, whereas HPV-unrelated p16+ tumors exhibited clinicodemographic profiles more similar to p16− tumors (Fig. 1C, Table S1). Regarding tumor location, both HPV-unrelated p16+ and p16− tumors arose significantly less frequently from the lateral wall compared with HPV-related tumors. In TNM classification, HPV-unrelated p16+ and p16− tumors less frequently exhibited advanced nodal involvement relative to primary tumor stage, suggesting similar clinical behavior between these groups (Fig. 1C). Survival analysis showed that p16⁻ tumors were associated with the poorest prognosis, whereas HPV-related tumors had significantly better outcomes. HPV-unrelated p16+ tumors displayed an intermediate prognosis between these two groups (Fig. 1D). In summary, a substantial subset of HPV-unrelated OPCs exhibited false-positive p16 expression and shared clinical features with p16− OPC, with an intermediate prognosis between HPV-related and p16− disease.

### HPV-unrelated p16-positive OPCs transcriptomically resemble p16-negative rather than HPV-related tumors

To investigate gene expression patterns associated with HPV infection and p16 expression status, RNA sequencing was performed on frozen biopsy specimens from 40 OPCs, including 17 HPV-related, 9 HPV-unrelated p16+, and 14 p16− tumors, together with 6 normal pharyngeal biopsy samples. Hierarchical clustering and heatmap analyses revealed two major groups: one comprising p16− and HPV-unrelated p16+ OPCs, and the other containing HPV-related OPCs and normal tissues (Fig. 2A). HPV-related OPCs formed a distinct subcluster separate from normal tissues, whereas p16− and HPV-unrelated p16+ OPCs remained intermixed without clear segregation. Principal component analysis showed a similar pattern, with HPV-related OPCs clustering closer to normal tissues, while HPV-unrelated p16+ and p16− OPCs clustered together in a distinct and overlapping group separate from HPV-related OPCs (Fig. 2B). These findings indicate that HPV-unrelated OPCs exhibit gene expression profiles distinct from those of HPV-related OPCs, irrespective of p16 expression status.

**Figure 2.**
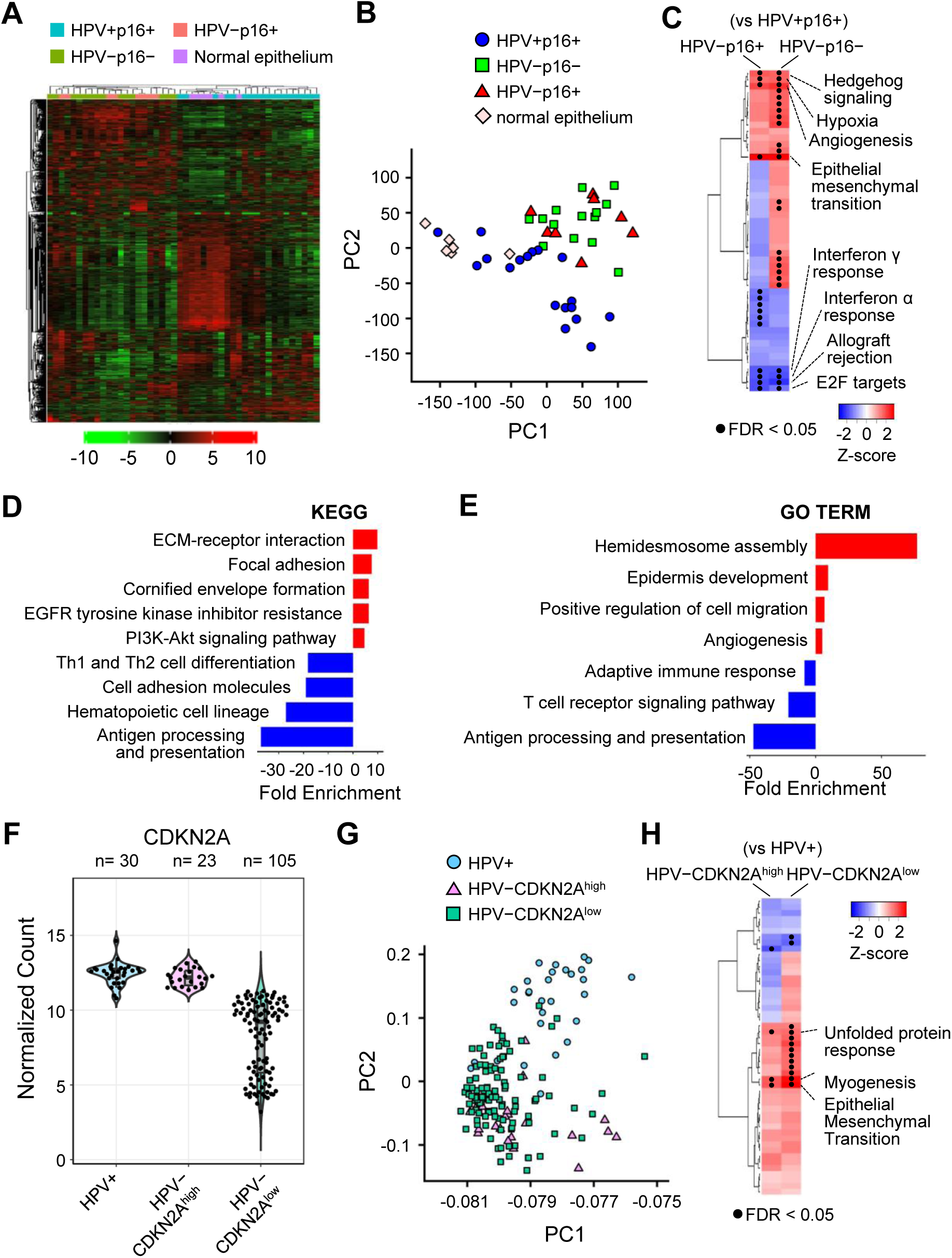
Transcriptome analysis of OPC stratified with HPV/p16 status. (A, B) Hierarchical clustering (A) and principal component analysis (B) of transcriptome data from our cohort. (C) GSEA of HPV−/p16+ and HPV−/p16− tumors, using HPV+/p16+ tumors as the reference group. Hallmark gene sets were analyzed. (D, E) KEGG pathway (D) and Gene Ontology (E) analyses showing pathways commonly altered in HPV-unrelated cancers compared with HPV-related cancers, regardless of p16 status. (F) Normalized CDKN2A transcript counts in the TCGA dataset. Based on CDKN2A expression levels covering 95% of HPV-related cancers, HPV-unrelated cancers were categorized as CDKN2A-high or CDKN2A-low. (G) Principal component analysis of TCGA cases. (H) GSEA of HPV−/CDKN2A-high and HPV−/CDKN2A-low tumors, using HPV+ tumors as the reference group.

We next assessed HPV-associated expression differences. Gene set enrichment analysis (GSEA) comparing HPV-unrelated p16+ and p16− OPCs with HPV-related tumors showed enrichment of invasion-associated programs, such as epithelial–mesenchymal transition (EMT) and angiogenesis, in HPV-unrelated tumors, whereas immune-response signatures (including interferon-α/γ signaling) and E2F targets were enriched in HPV-related tumors (Fig. 2C, Fig. S1A, Tables S2, S3). Differential expression analysis identified 257 differentially expressed genes (DEGs) consistently upregulated and 75 DEGs consistently downregulated in both HPV-unrelated groups versus HPV-related tumors (Fig. S1B, Tables S4, S5). Consistent with the GSEA results, pathway and gene ontology analyses of these shared DEGs indicated upregulation of genes related to cell adhesion and migration and downregulation of immune-related genes in HPV-unrelated tumors (Fig. 2D, E, Tables S6, S7). These findings suggest that, regardless of p16 status, HPV-unrelated OPCs exhibit a more invasive transcriptional program and weaker immune activation than HPV-related OPCs.

To validate our findings, we analyzed RNA sequencing data from The Cancer Genome Atlas (TCGA). Among 158 oral and oropharyngeal cancers, HPV transcripts were detected in 30 cases (19.0%), defining HPV-related tumors (11). Because p16 IHC is not consistently annotated in TCGA, we used CDKN2A transcript abundance (encoding p16) as a surrogate, supported by its strong correlation with p16 immunostaining in our cohort (Fig. S2). As in our samples, high CDKN2A expression was observed not only in most HPV-related tumors (28/30, 93.3%) but also in a subset of HPV-unrelated tumors (23/128, 18.0%) (Fig. 2F), indicating appreciable p16/CDKN2A overexpression in HPV-unrelated OPCs.

We then compared HPV-related tumors with HPV-unrelated tumors stratified by CDKN2A expression in the TCGA data. HPV-unrelated CDKN2A-high tumors had significantly better outcomes than CDKN2A-low tumors (Fig. S3A). Unsupervised clustering and principal component analysis separated HPV-related from HPV-unrelated tumors, whereas CDKN2A-high and -low cases were intermixed within the HPV-unrelated group (Figs. 2G, S3B). GSEA showed that HPV-unrelated tumors were enriched for EMT signatures regardless of CDKN2A status (Fig. 2H, S3C, Tables S8, S9). The 840 DEGs upregulated in HPV-unrelated cancers were enriched for EMT signatures, whereas the 1,066 downregulated DEGs were associated with immune signatures (Fig. S3D, Table S10, S11, S12). Taken together, both our cohort and TCGA indicate that HPV-unrelated OPCs with p16/CDKN2A overexpression have a better prognosis than p16/CDKN2A-low tumors, yet retain HPV-unrelated expression profiles characterized by increased invasiveness and reduced immune activity compared with HPV-related OPCs.

### CDKN2A/p16 is the only consistent transcriptomic difference between HPV-unrelated CDKN2A/p16-high and -low tumors

We further compared gene expression between HPV-unrelated CDKN2A/p16-positive and - negative cancers. In our cohort, 19 genes, including CDKN2A, were significantly upregulated in HPV-unrelated p16+ versus p16− OPCs (Fig. 3A, Table S13), whereas TCGA showed 254 upregulated genes in HPV-unrelated CDKN2A-high versus-low tumors (Fig. 3B, Table S14). Across both datasets, however, CDKN2A was the only consistently upregulated gene, and no genes were consistently downregulated (Fig. 3C). GSEA further showed that immune- and inflammation-related pathways were commonly downregulated in CDKN2A/p16-positive cases, while no gene sets consistently upregulated across both datasets (Fig. 3D, 3E, Tables S15, S16). Together with the intermixed clustering of CDKN2A/p16-positive and -negative tumors (Fig. 2A, Fig. S3B), these results indicate minimal reproducible transcriptomic differences beyond CDKN2A/p16 itself, suggesting that the better prognosis of HPV-unrelated CDKN2A/p16-positive cancers compared with their CDKN2A/p16-negative counterparts may be largely attributable to CDKN2A/p16 expression.

**Figure 3.**
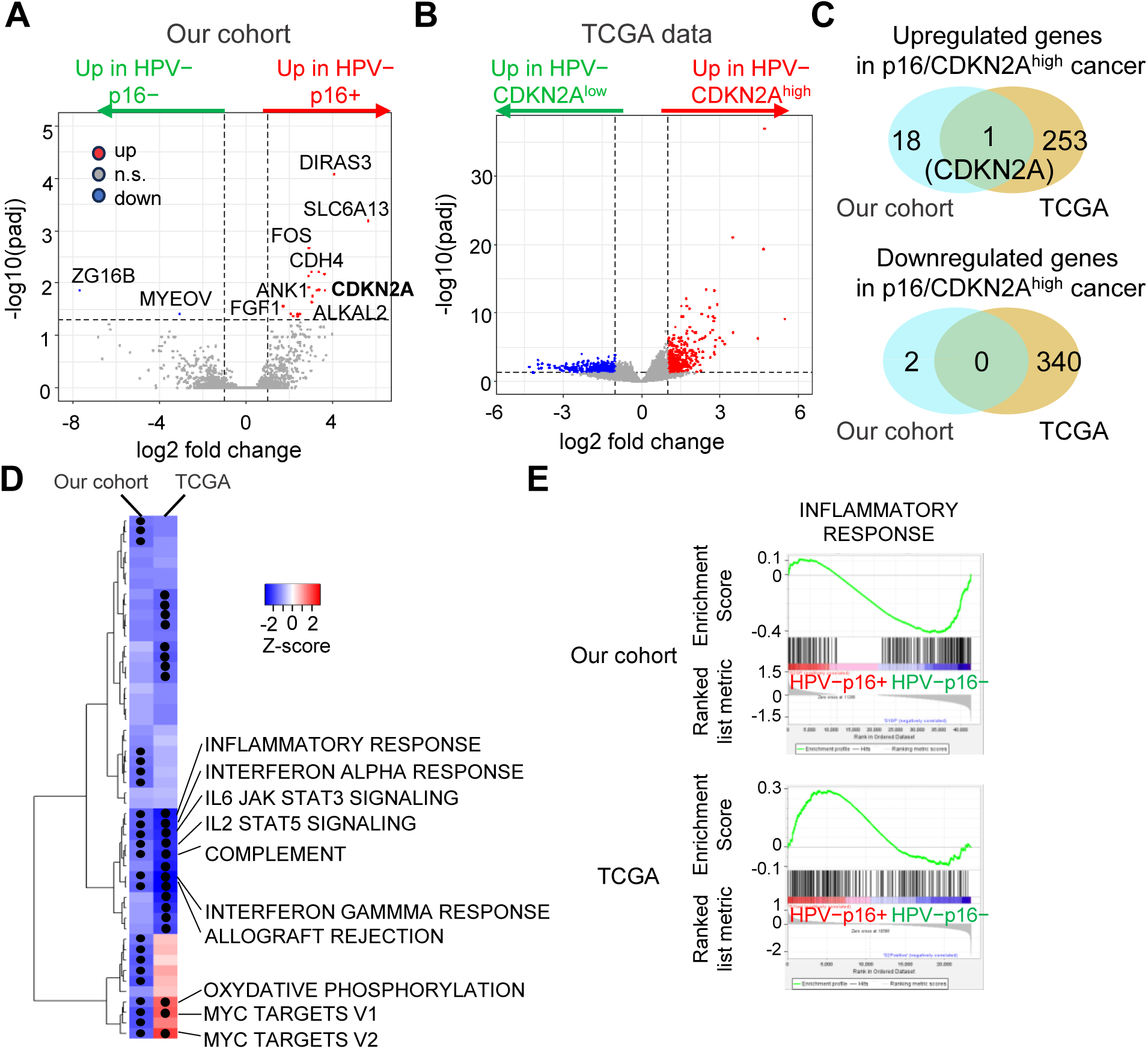
Comprehensive gene expression differences between p16/CDKN2A-high and p16/CDKN2A-low HPV-unrelated OPC. (A) Volcano plot of differentially expressed genes between HPV−/p16+ and HPV−/p16− OPCs in our cohort. (B) Volcano plot of differentially expressed genes between HPV−/CDKN2A-high and HPV−/CDKN2A-low OPCs in the TCGA dataset. (C) Venn diagrams showing the numbers of genes significantly upregulated or downregulated in p16/CDKN2A-high versus p16/CDKN2A-low HPV-unrelated OPCs across our cohort and TCGA data. (D, E) GSEA comparing HPV− p16/CDKN2A-high and HPV− p16/CDKN2A-low tumors.

### CDK6 overexpression distinguishes HPV-unrelated OPCs from HPV-related cancers, irrespective of CDKN2A/p16 expression status

Given that HPV-related and HPV-unrelated OPCs show globally distinct expression profiles regardless of p16 status, we sought genes that correlate more strongly with HPV status than p16. Across both our cohort and the TCGA dataset, we identified 146 genes consistently upregulated and 51 genes consistently downregulated in HPV-unrelated cancers relative to HPV-related cancers (Fig. S4A, Tables S17). Among the genes upregulated in HPV-unrelated cancers, 12 genes were annotated to the KEGG “Pathways in cancer” pathway, including EMT-related genes (COL17A1, COL4A5, COL4A6, ITGA6, LAMA3, LAMC2, and MMP1), Hedgehog signaling genes (GLI1 and GLI2), EGFR pathway genes (EGFR and TGFA), and CDK6 (Fig. 4A, S4B). Importantly, CDK6 is a cell-cycle kinase functionally inhibited by p16 (Fig. 4B). Both in our cohort and the TCGA dataset, CDK6 was overexpressed in HPV-unrelated cancers compared with HPV-related cancers irrespective of CDKN2A status, a pattern that was distinctive among other Rb-pathway genes (Fig. 4C, S4B, S4C). Consistently, IHC showed strong CDK6 protein expression in the majority of HPV-unrelated OPCs regardless of p16 status, whereas most HPV-related OPCs showed low or undetectable CDK6 (Fig. 4D, 4E). In contrast, CDK4, another p16 target, did not differ between HPV-related and HPV-unrelated tumors at the protein level (Fig. S4D). These findings indicate that CDK6 overexpression is a defining feature of HPV-unrelated OPCs independent of p16 status, and underscore the central role of the p16–CDK6 axis in HPV-unrelated OPC carcinogenesis.

**Figure 4.**
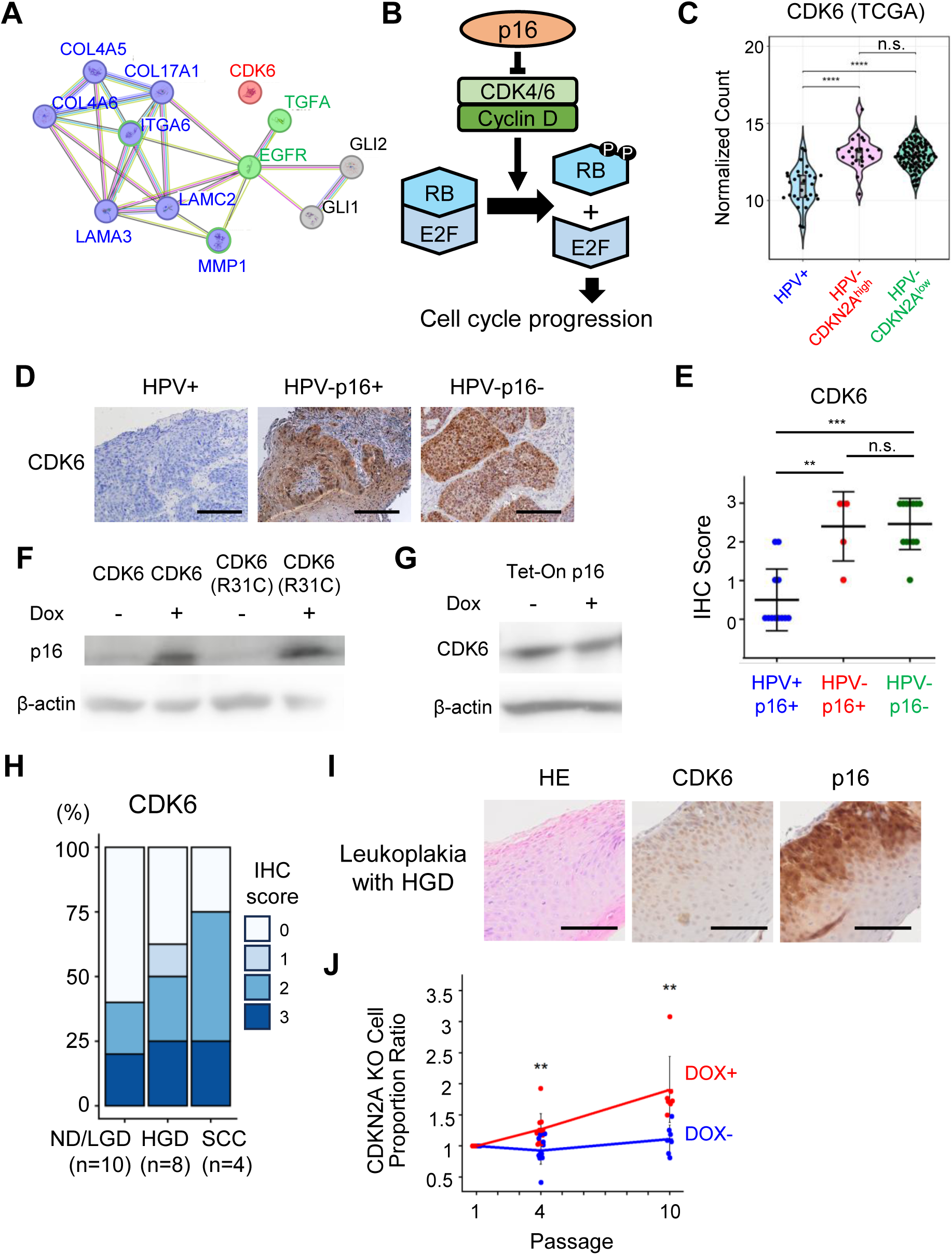
CDK6 overexpression in HPV-unrelated OPC. (A) STRING network visualization of genes in the “Pathways in cancer” pathway that were commonly upregulated in HPV-unrelated OPC. (B) Schematic diagram of the p16–RB pathway. (C) CDK6 expression in RNA sequencing data stratified by HPV and CDKN2A status in the TCGA dataset. (D) Representative immunohistochemistry images of CDK6 staining. Scale bar, 200 μm. (E) IHC scores for CDK6 staining. (F) Western blot showing p16 expression in RPE1 cells with doxycycline-inducible CDK6 expression. (G) Western blot showing CDK6 expression following p16 transduction in CDKN2A-deficient FaDu cells. (H) Proportion of CDK6 expression levels in leukoplakia. CDK6 immunohistochemical staining was evaluated using a scoring system, and the distribution of scores is shown by histopathological grade. ND/LGD, non-dysplastic/low-grade dysplasia; HGD, high-grade dysplasia; SCC, squamous cell carcinoma in situ or carcinoma with minimal invasion. (I) HE and immunohistochemical images of oral leukoplakia with high-grade dysplasia showing p16 expression. Scale bars, 50 μm. (J) Clonal expansion of CDKN2A-deficient cells in CDK6-overexpressing RPE1 cells. The fraction of CDKN2A indels was quantified over time in RPE1 cells expressing the CDK6 (R31C) mutant in a doxycycline-dependent manner. DOX, doxycycline. Statistical analyses were performed using the Mann–Whitney U test in (C) and (E), and Welch’s t-test in (J). **p < 0.01, ***p < 0.001. n.s., not significant.

### Aberrant expression of the p16–CDK6 axis in HPV-unrelated carcinogenesis

We next investigated potential mechanisms underlying aberrant p16 expression during HPV-unrelated OPC carcinogenesis. Because CDK6 promotes cell-cycle progression and is broadly overexpressed in HPV-unrelated OPCs, we hypothesized that CDK6 upregulation occurs first and subsequently induces p16 via feedback. To examine this, we induced CDK6 expression in the non-transformed human epithelial cell line RPE1 in a doxycycline-dependent manner and found that CDK6 induction increased p16 expression (Fig. 4F). Conversely, p16 induction caused cell-cycle arrest and prevented the upregulation of CDK6 (Fig. 4G). Furthermore, immunohistochemical assessment of CDK6 expression in oral leukoplakia, a premalignant lesion of oral cancer, frequently revealed CDK6 overexpression in high-grade dysplasia and in early squamous cell carcinoma (SCC) lesions (SCC in situ or minimally invasive SCC) (Fig. 4H, I). Among the eight cases of high-grade dysplasia and four cases of early SCC, p16 positivity was observed only in two high-grade dysplasia cases: one was HPV-positive without concomitant CDK6 overexpression, whereas the other was HPV-negative with CDK6 overexpression (Fig. 4I). Despite the limited sample size and the anatomical difference between OPC and leukoplakia/oral cancer, these findings suggest that CDK6 overexpression is a frequent early event in the development of head and neck SCC and can drive p16 overexpression.

Because p16 is a tumor suppressor that restrains cell proliferation, loss of CDKN2A, which encodes p16, can act as a key driver of carcinogenic evolution. We further hypothesized that selective pressure for CDKN2A loss is imposed specifically under conditions that provoke feedback p16 upregulation, and that CDK6 overactivation may provide such a selective context. To test this idea, we tracked the clonal dynamics of CDKN2A-deficient cells in our doxycycline-inducible CDK6 system in RPE-1 cells. We first introduced CRISPR-mediated CDKN2A disruptions to generate a mixed population containing ∼40% CDKN2A-knockout cells. Notably, in the absence of doxycycline, CDKN2A-knockout cells showed no appreciable clonal expansion. In contrast, doxycycline-induced CDK6 activation resulted in a marked expansion of CDKN2A-knockout clones (Fig. 4J). Consistent with this clonal replacement of CDKN2A-deficient cells, we also identified a case that was initially diagnosed as a p16–high, HPV-unrelated carcinoma but became negative for p16 staining during follow-up (Fig. S5). Taken together, these results suggest that CDK6 activation elicits a p16-mediated antiproliferative feedback, and that this feedback state in turn imposes selective pressure for CDKN2A loss, promoting clonal outgrowth of CDKN2A-deficient cells.

### CDKN2A frameshifts can yield false-positive p16 IHC via a chimeric p14ARF

We next investigated genetic alterations in cancer-related genes, including components of the p16–CDK6 axis, in HPV-unrelated OPCs using targeted capture sequencing. Homozygous CDKN2A deletions were observed exclusively and frequently in p16⁻ OPCs (6/8, 75%), whereas loss of heterozygosity of the CDKN2A gene was detected in all other cases, including p16⁺ OPCs (Fig. 5). In addition, nucleotide-level alterations, particularly indels, were frequently detected in HPV-unrelated p16⁺ tumors (Fig. 5). Among other genes in the Rb pathway, nucleotide alterations were detected exclusively in RB1, and only in two HPV-unrelated p16⁺ cases (Fig. 5). Although RB1 disruption can also contribute to feedback upregulation of CDKN2A (18), its low frequency in HPV-unrelated p16⁺ OPC suggests that CDK6 upregulation is the primary driver of high p16 expression in this subgroup.

**Figure 5.**
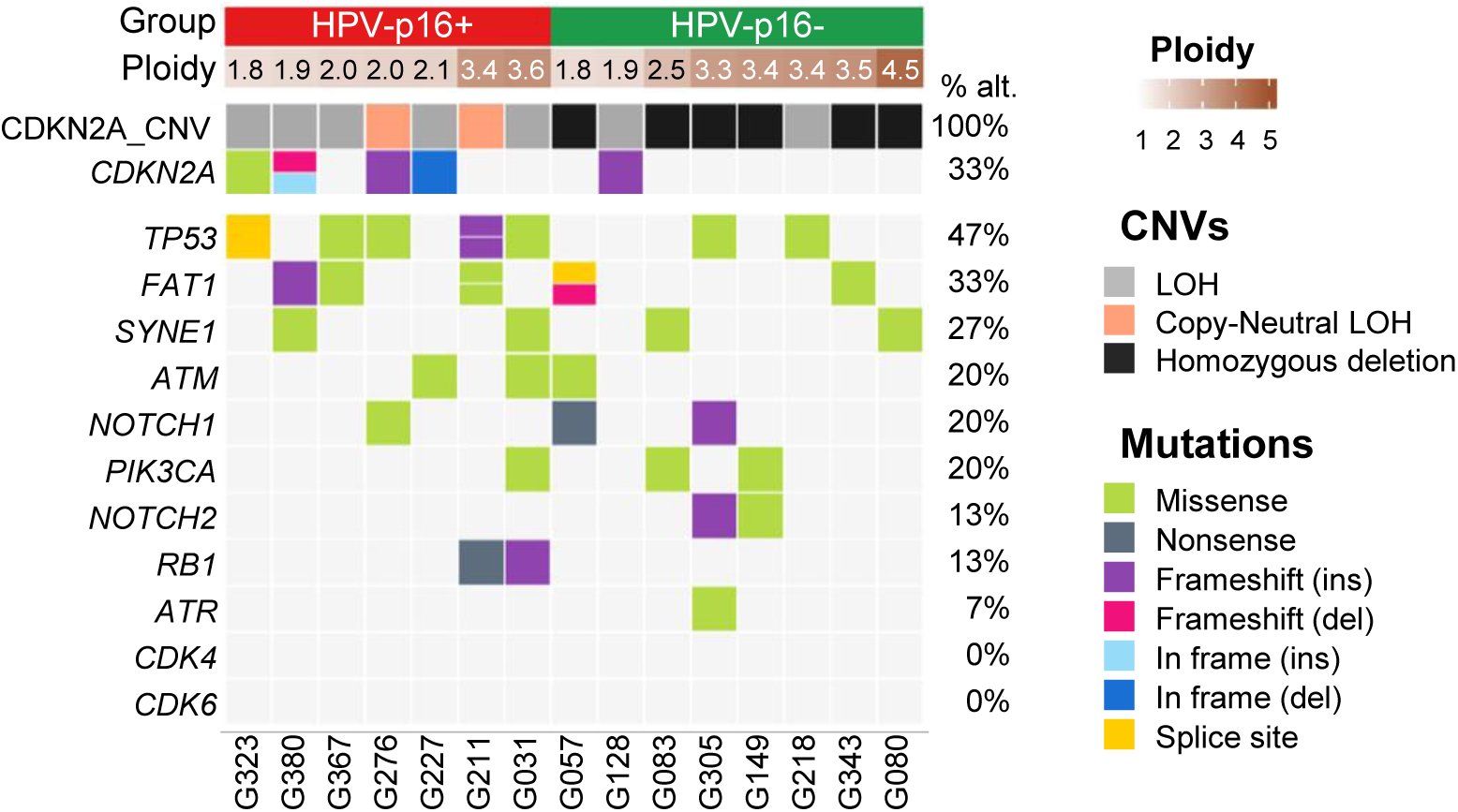
Oncopanel analysis of HPV-unrelated OPC. Ploidy and detected nucleotide alterations in major cancer-associated genes are shown for each case. For CDKN2A, copy number alterations are also presented. The percentages represent the frequency of cases with each alteration among all analyzed cases.

Because CDKN2A is minimally expressed in normal cells, the CDKN2A transcripts detected by transcriptome analysis were almost exclusively tumor-derived, allowing us to corroborate tumor CDKN2A nucleotide alterations at the RNA level (Fig. S6). By integrating capture sequencing and transcriptome data, we identified five tumors with nucleotide alterations in CDKN2A accompanied by complete loss of the wild-type allele (Fig. 6A, B). Notably, two of these tumors were scored as p16-positive by IHC despite expressing only frameshift transcripts predicted to truncate p16 (Fig. 6B). Importantly, the p16 antibody clone E6H4 recognizes an epitope C-terminal to the frameshift sites in these tumors (Fig. S7A) (19, 20) and therefore should not detect a truncated p16 product. Consistent with this, ectopic expression of cohort-derived CDKN2A variants in p16-deficient FaDu cells showed that in-frame mutant p16 was readily detected, whereas frameshifted p16 was not detectable by immunostaining (Fig. S7B).

**Figure 6.**
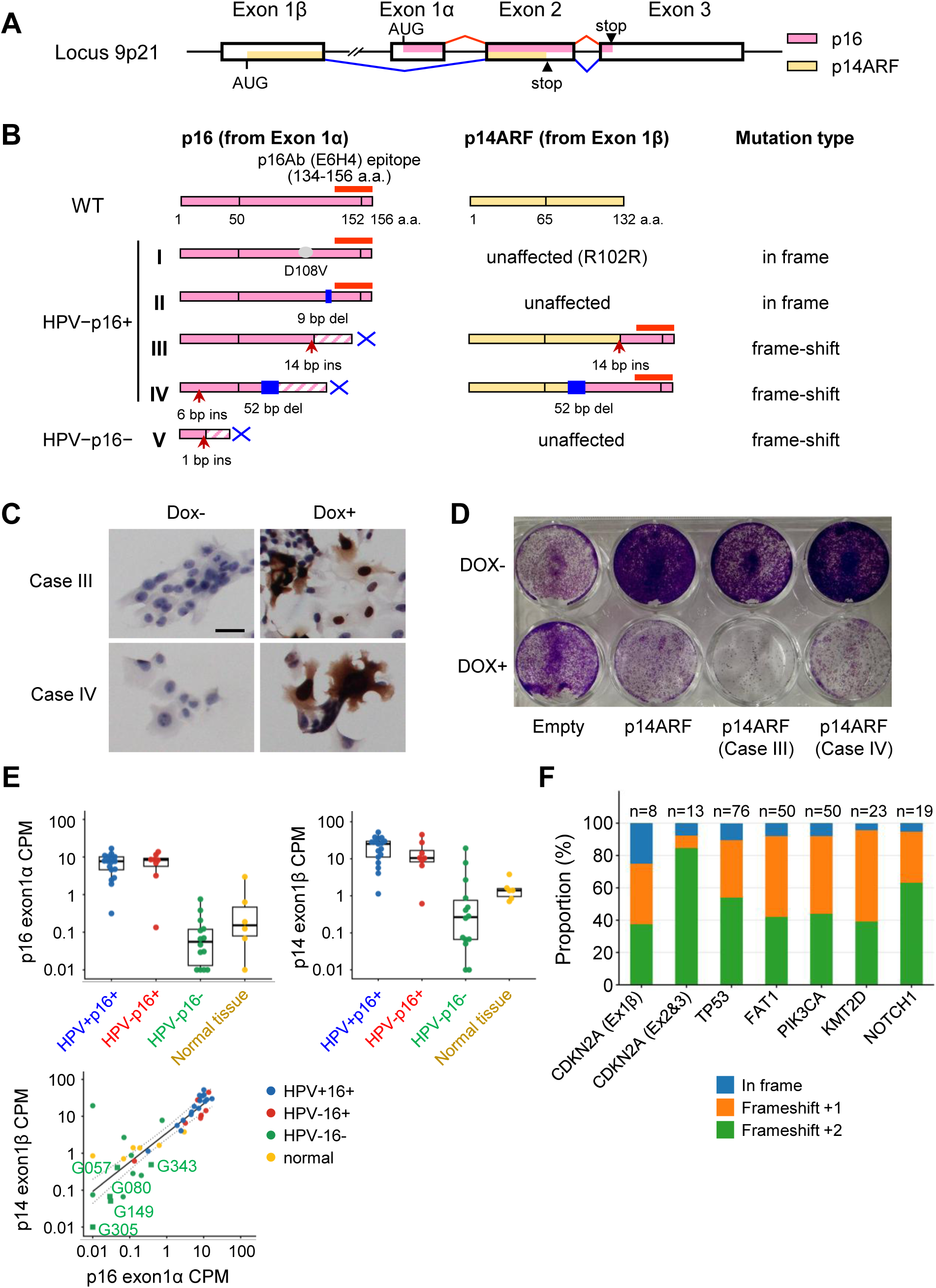
Impact of frameshift mutations in CDKN2A locus. (A) Schematic representation of p16 and p14ARF encoded at the CDKN2A locus. Pink and yellow indicate the amino acid-coding regions of p16 and p14ARF, respectively. (B) Detected nucleotide alterations in the CDKN2A locus and their predicted effects on the p16 and p14ARF proteins. The red bar indicates the location of the epitope recognized by the anti-p16 antibody (clone E6H4). (C) Detection of chimeric p14ARF proteins using an anti-p16 antibody (clone E6H4). The chimeric proteins were expressed in CDKN2A-deficient FaDu cells and analyzed by immunostaining. Scale bars, 20 μm. (D) Crystal violet staining showing the effects of chimeric p14ARF proteins on cell proliferation. The chimeric proteins were expressed in FaDu cells in a doxycycline-dependent manner, and crystal violet staining was performed 7 days later. (E) Correlation between p16 and p14ARF expression levels in our transcriptome data. (F) Frequencies of frameshift mutations in cancer-related genes stratified by the resulting reading-frame shift in TCGA head and neck cancer samples.

To explain why p16 IHC remained positive in the two tumors harboring CDKN2A-truncating frameshift indels, we considered the dual-coding architecture of the CDKN2A locus. CDKN2A produces two distinct proteins, p16 and p14ARF, via alternative first exons and reading frames; thus, these frameshift variants could affect not only p16 but also p14ARF (Fig. 6B). In silico analysis predicted that the variants would remodel the C-terminus of p14ARF to a sequence matching the C-terminal region of canonical p16 (Fig. 6B). Consistent with this, ectopic expression of the aberrant p14ARF in FaDu cells resulted in strong staining with the p16 (E6H4) antibody (Fig. 6C) and demonstrated functional activity in suppressing tumor cell proliferation (Fig. 6D). Furthermore, our transcriptome analysis focusing on exon 1α and exon 1β, which are specific to p16 and p14ARF, respectively, showed a strong correlation between p16 and p14ARF transcript levels regardless of HPV status, suggesting that p14ARF is also highly expressed in p16-positive cancers (Fig. 6E). In the TCGA head and neck cancer cohort, +2 frameshift mutations that truncate p16 and generate a chimeric protein fusing the N terminus of p14ARF to the C terminus of p16 were frequently observed in CDKN2A exons 2 and 3, which are shared by p16 and p14ARF (Fig. 6F). Together, these findings suggest that a subset of HPV-unrelated OPCs are p16 IHC positive despite harboring p16 truncating frameshifts, although why such frameshifts recur in CDKN2A remains unclear.

## Discussion

Our analysis of more than 350 evaluable cases confirmed the high sensitivity of p16 IHC for identifying oncogenic HPV-related tumors. However, p16 positivity did not always reflect true oncogenic HPV infection, resulting in a specificity of only 88%. This performance is consistent with previous reports demonstrating that a subset of HPV-unrelated OPCs are p16-positive by IHC (i.e., false-positive for HPV status) (13, 21). Notably, HPV-unrelated p16-positive OPCs closely resembled p16-negative tumors in their global transcriptomic profiles and inferred etiologic and progression patterns, supporting the interpretation that HPV-unrelated p16-positive OPC represents a distinct subgroup within HPV-unrelated disease rather than reflecting false-negative HPV-PCR. Indeed, in transcriptome comparisons within HPV-unrelated OPC between tumors with and without high p16 (or CDKN2A) expression, the only reproducible differential expression across both our cohort and TCGA was upregulation of CDKN2A itself. Despite this remarkably limited transcriptomic separation, HPV-unrelated p16-positive tumors showed a more favorable prognosis than p16-negative tumors, suggesting that the biological impact of p16 expression may not be fully captured by bulk transcriptome analyses. Although this survival difference did not reach statistical significance in our cohort, likely owing to the limited sample size and differences in baseline clinicopathological characteristics between the groups, it was significant in the independent TCGA cohort and is consistent with findings from a recent large multicenter study (13). Importantly, beyond its canonical role as an inhibitor of CDK4/6-mediated cell-cycle progression, p16 has been implicated in diverse non-canonical functions, including regulation of nucleotide metabolism and the DNA damage response (22, 23), and these activities have recently been linked to therapeutic responsiveness in HPV-related head and neck SCC (23). These observations raise the possibility that non-canonical functions of p16 contribute to the favorable prognosis of HPV-unrelated p16-positive OPC, supporting the clinical value of p16 expression beyond its role as a surrogate marker of HPV infection.

Our transcriptome analyses further recapitulated key biological differences between HPV-related and HPV-unrelated OPC. HPV-related tumors showed E2F target activation and increased expression of immune-related genes, whereas HPV-unrelated tumors preferentially upregulated programs related to hypoxia, angiogenesis, and EMT. These findings align well with previous reports demonstrating stronger immune activation in HPV-driven OPC and a more aggressive biology in HPV-unrelated disease (5, 24). The observed E2F activation in HPV-related OPC is also mechanistically consistent with HPV E7–mediated Rb inactivation, which releases E2F from Rb-dependent repression (25). Beyond these differences, we identified CDK6 upregulation as a recurrent feature of HPV-unrelated OPC regardless of p16 status. Functional inactivation of Rb by HPV E7 may reduce the dependence of HPV-related tumors on CDK6 activity, potentially contributing to the relatively low CDK6 expression observed in these tumors. Accordingly, combined assessment of p16 and CDK6 may improve discrimination of HPV-related tumors beyond p16 IHC alone. Although CDK6 overexpression in human cancers is frequently driven by gene amplification (26), no evidence of CDK6 amplification was observed in our cohort (data not shown). Alternative mechanisms, including epigenetic dysregulation and disruption of p16-mediated negative feedback following CDKN2A loss, have also been reported to increase CDK6 expression and may account for a subset of HPV-unrelated OPCs (27). However, the latter mechanism is unlikely to explain the frequent CDK6 upregulation observed in p16-positive HPV-unrelated tumors. Although the mechanisms underlying CDK6 upregulation remain unclear, our findings suggest that CDK6 may play a more prominent role than CDK4 in HPV-unrelated OPC despite the largely overlapping functions of these p16-targeted cyclin-dependent kinases. Clarifying the mechanisms responsible for CDK6 upregulation and why CDK6 is preferentially engaged over CDK4 in HPV-unrelated OPC will be an important objective of future studies. Although this proposed evolutionary model remains to be validated in larger longitudinal studies, several observations from our study support this hypothesis. Consistent with an early role for CDK6 activation during oral/oropharyngeal tumorigenesis, leukoplakia with high-grade dysplasia frequently exhibited CDK6 overexpression, with a subset also showing p16 overexpression (Fig. 4H, 4I). Furthermore, HPV-unrelated p16-positive OPCs frequently harbored LOH and CDKN2A nucleotide alterations, whereas HPV-unrelated p16-negative tumors predominantly exhibited homozygous CDKN2A deletion. Together, these findings are compatible with progressive CDKN2A inactivation during HPV-unrelated OPC development and support a model in which at least a subset of tumors evolve from a CDK6-high/p16-high state toward a CDKN2A-inactivated state.

Given the selective advantage conferred by CDKN2A loss under CDK6-driven pressure and the favorable prognosis associated with p16 positivity in HPV-unrelated OPC, p16 upregulation appears to restrain tumor progression through its canonical and/or non-canonical functions. Consistent with selective pressure to overcome this restraint, HPV-unrelated p16-positive tumors frequently exhibited LOH or nucleotide alterations affecting at least one CDKN2A allele. Strikingly, 2 of 7 (28.6%) HPV-unrelated p16-positive OPCs remained positive by p16 IHC despite complete loss of the wild-type CDKN2A allele and expression exclusively of frameshift-indel CDKN2A transcripts predicted to truncate p16. These alterations are predicted to generate a chimeric p14ARF protein with a p16-like C-terminus, resulting in immunoreactivity with the commonly used E6H4 antibody. Although the functional consequences of this C-terminal alteration remain uncertain, our in vitro assays suggest that the chimeric protein retains growth-inhibitory activity comparable to wild-type p14ARF. Although such p14ARF–p16 chimeric proteins are likely uncommon in OPC overall, they represent an important diagnostic pitfall for HPV status assessment because they can produce positive p16 immunostaining despite complete functional loss of wild-type p16. In summary, our findings underscore the limitations of relying on p16 IHC alone for molecular classification of OPC. By focusing on HPV-unrelated tumors with p16 overexpression, we support the concept that dysregulation of the p16–CDK6 axis is an important contributor to HPV-unrelated OPC tumorigenesis. CDK6 overexpression has been implicated in resistance to CDK4/6 inhibitors (28, 29), which are currently being investigated as a therapeutic strategy for OPC (30, 31). Prospective studies tracking alterations of the p16–CDK6 axis during the evolution of HPV-unrelated OPC should refine diagnostic algorithms, improve biological stratification, and facilitate the development of biomarker-guided therapeutic strategies.

## Abbreviations

GSEA: Gene set enrichment analysis
HPV: human papillomavirus
IHC: immunohistochemistry
OPC: oropharyngeal squamous cell carcinoma
SCC: squamous cell carcinoma

## Acknowledgements

We thank the Core Instrumentation Facility, Research Institute for Microbial Diseases, Osaka University, for technical assistance with FACS sorting, and the Genome Information Research Center, Osaka University, for conducting RNA sequencing.

## Data availability

The RNA sequencing data generated in this study have been deposited in the DDBJ Sequence Read Archive (https://www.ddbj.nig.ac.jp) under accession number DRA029340. The authors declare that all other data are available upon request.

## Materials and methods

### Patients and assessment of HPV and p16 status

A total of 352 patients with oropharyngeal cancer (OPC) who were treated at The University of Osaka Hospital or Osaka International Cancer Institute between July 1, 2015, and June 30, 2019, were included in this study. HPV and p16 status were assessed using tumor specimens obtained at initial presentation. IHC staining for p16 was performed on formalin-fixed, paraffin-embedded tissue sections using the p16 antibody clone E6H4 (Roche, Basel, Switzerland) as part of routine clinical practice covered by the Japanese national health insurance system. Staining was conducted according to standard diagnostic procedures, and tumors showing p16 expression in ≥70% of tumor cells were classified as p16-positive. For HPV testing, DNA was extracted from frozen tumor specimens collected at the initial visit using the QIAamp DNA Mini Kit (Qiagen, Hilden, Germany) or the DNeasy Blood & Tissue Kit (Qiagen). HPV detection and genotyping were performed using nested polymerase chain reaction followed by direct sequencing of the amplified products, as previously described (32). Clinical data were obtained from patients’ medical records. This study was conducted in accordance with the Declaration of Helsinki and was approved by the Ethics Committee of The University of Osaka Hospital (approval no. 22125-3). Patient participation was based on either written informed consent or an ethics committee–approved opt-out procedure, as appropriate for each component of the study.

### Cell culture

The human retinal pigment epithelium cell line hTERT-RPE1 (RPE1) was obtained from Lonza (Basel, Switzerland). The human hypopharyngeal carcinoma cell line FaDu was obtained from the American Type Culture Collection (ATCC; Manassas, VA, USA). All cells were cultured in Dulbecco’s modified Eagle’s medium (DMEM; Nacalai Tesque, Kyoto, Japan) supplemented with 10% fetal bovine serum (Nichirei Bio Sciences, Tokyo, Japan). All cell lines tested negative for mycoplasma contamination.

RPE1 cell lines with tetracycline-inducible CDK6 expression were generated using the Sleeping Beauty transposon system, as previously described (33). A Sleeping Beauty transposon plasmid encoding CDK6 or a constitutively active CDK6 mutant (R31C) was constructed by subcloning PCR-amplified CDK6 cDNA (wild-type or R31C) into pSBtet-RP (Addgene plasmid #60497) (34). The RFP cassette in pSBtet-RP was deleted, and an IRES–emiRFP670 cassette was inserted downstream of CDK6. RPE1 cells were co-transfected with the modified transposon vector and pCMV(CAT)T7-SB100 (Addgene plasmid #34879) (35), which encodes the SB100X transposase. Cells with stable integration were selected with puromycin for more than 1 month.

CDKN2A knockout (KO) cells were generated using the CRISPR–Cas9 system. Cas9 Nuclease V3 (Integrated DNA Technologies, IDT), tracrRNA (IDT), and a CDKN2A-targeting crRNA (CCCAACGCACCGAATAGTTA; IDT) were assembled and delivered into cells using Lipofectamine CRISPRMAX (Thermo Fisher Scientific). The initial KO efficiency was approximately 40%, and changes in KO efficiency were subsequently monitored over time. To quantify KO efficiency, the CRISPR target region was PCR-amplified using primers CTTTGCTATTTTGCCCGTGCC and CATCTATGCGGGCATGGTTACT, and the amplicons were gel-purified and subjected to Sanger sequencing (Central Instrumentation Laboratory, Research Institute for Microbial Diseases, The University of Osaka). Indel frequencies were estimated from the sequencing chromatograms using ICE (Inference of CRISPR Edits) analysis pipeline (Synthego, Redwood City, CA, USA).

### RNA sequencing analysis

Total RNA was extracted from fresh-frozen biopsy specimens using the RNeasy Micro Kit (Qiagen) according to the manufacturer’s instructions. Poly(A)+ RNA was isolated using the NEBNext Poly(A) mRNA Magnetic Isolation Module (New England Biolabs, Ipswich, MA, USA), and strand-specific sequencing libraries were prepared with the NEBNext Ultra II Directional RNA Library Prep Kit (New England Biolabs). Paired-end sequencing (2 × 150 bp) was performed on the Illumina NovaSeq 6000 platform (Illumina, San Diego, CA, USA) at Rhelixa, Inc. (Tokyo, Japan), generating approximately 40 million reads (approximately 6 Gb of sequence data) per sample. Sequence reads were aligned to the human hg38 reference genome using STAR (version 2.7.10b) (36). The reads per gene were counted using HTSeq (version 2.0.2) (37). Differential gene expression analysis was performed using edgeR (version 3.36.0) (38) in R, and genes with an adjusted P value < 0.05 were considered significantly differentially expressed. For visualization purposes, expression levels were calculated as Trimmed Mean of M-values (TMM)-normalized log2 counts per million (CPM) using edgeR and are presented in the figures. Gene set enrichment analysis was performed by the Broad Institute GSEA (version 4.3.3) (39). KEGG pathway and Gene Ontology enrichment analyses of DEGs were performed using the Database for Annotation, Visualization and Integrated Discovery (DAVID, version 2021) (40, 41). Sequence alignments and genomic alterations were visually inspected using the Integrative Genomics Viewer (IGV, version 2.19.7)(42).

### Immunostaining and histology

Formalin-fixed, paraffin-embedded tissue specimens were sectioned at 5 µm and subjected to IHC staining. Primary antibodies against CDK6 (ab124821, Abcam, Cambridge, UK), CDK4 (sc-260, Santa Cruz, Dallas, TX, USA), Ki67 (ab15580, Abcam), and p16 (clone E6H4; CINtec Histology Kit-9511, Roche, Basel, Switzerland) were used. Detection was performed using anti-rabbit (MP-7401, Vector Laboratories, Burlingame, CA, USA) or anti-mouse (MP-7402, Vector Laboratories) secondary antibodies, followed by visualization with ImmPACT DAB (SK-4105, Vector Laboratories). All sections were incubated with primary antibodies overnight at 4°C. Stained slides were imaged using a BX53 upright microscope or a VS200 Research Slide Scanner (EVIDENT, Tokyo, Japan) and analyzed with OlyVIA v4.1 software (EVIDENT).

### IHC scoring

IHC staining was scored as follows: 3+, strong staining of both the nucleus and cytoplasm; 2+, staining of either the nucleus or cytoplasm, or staining intensity intermediate between 1+ and 3+; 1+, faint cytoplasmic staining regardless of nuclear staining; and 0, complete absence of staining in both the nucleus and cytoplasm. Cases in which no staining was observed in any cells, including lymphocytes, or in which staining could not be reliably evaluated for technical reasons were considered unevaluable. For analysis, cases with a score of 2+ or higher were classified as positive. For CDK6 IHC scoring, cases in which the surrounding lymphocytes did not show clear CDK6 immunoreactivity were considered unevaluable and excluded from the analysis.

### Transfection, immunocytochemistry and crystal violet staining

RPE1 and FaDu cells were transfected with plasmids using Lipofectamine 3000 (Thermo Fisher Scientific) according to the manufacturer’s instructions. Two to three days after transfection, cells were subjected to immunocytochemical analysis. For immunocytochemistry, cells were fixed with 4% paraformaldehyde, permeabilized, and stained using the same antibodies and staining procedures as those described for immunohistochemistry. Slides were photographed using an Olympus BX53 microscope (Evident Corporation, Tokyo, Japan). For crystal violet staining, culture medium was removed, and cells were washed with phosphate-buffered saline (PBS) and fixed with 4% paraformaldehyde for 15 min at room temperature. After two washes with PBS, cells were stained with 0.25% crystal violet solution for at least 10 min and then air-dried. For crystal violet staining, Cells were fixed with 4% paraformaldehyde and stained with 0.25% crystal violet as previously described (33).

### Western blot

Whole-cell lysates were prepared in RIPA buffer supplemented with a protease inhibitor cocktail (Nacalai Tesque). Protein concentrations were quantified using a Protein Assay Kit (Takara Bio). Equal amounts of protein were mixed with Laemmli sample buffer, and heated at 95°C for 5 min. The samples were then subjected to SDS–PAGE and transferred onto polyvinylidene difluoride membranes (EMD Millipore, Billerica, MA, USA). Membranes were blocked with 5% skim milk and incubated with primary antibodies against p16 (sc-56330, Santa Cruz Biotechnology) and β-actin (A5316, Sigma-Aldrich). After washing, membranes were probed with horseradish peroxidase-conjugated secondary antibodies (Cell Signaling Technology). Immunoreactive signals were detected using Amersham ECL Prime (Cytiva, Marlborough, MA, USA) and visualized with an ImageQuant 800 imaging system (Cytiva).

### Capture sequencing and data analysis

Genomic DNA was isolated from fresh-frozen biopsy specimens and peripheral blood samples using either the QIAamp DNA Mini Kit or the DNeasy Blood & Tissue Kit (Qiagen, Hilden, Germany) according to the manufacturer’s protocols. Sequencing libraries were generated with the Twist EF 2.0 Library Preparation Kit (Twist Biosciences, South San Francisco, CA, USA). Target enrichment was performed using the Biken Cancer Panel v2 (TE-94765031, Twist Biosciences) together with a CNV backbone panel (TE-98081791, Twist Biosciences). Captured libraries were sequenced as 100-bp paired-end reads on the MGI DNBSEQ-G400RS platform (MGI, Shenzhen, China) at the Genome Information Research Center, Research Institute for Microbial Diseases, The University of Osaka.

Somatic variant calling was performed using the Genome Analysis Toolkit (GATK, version 4.6.2.0) (43). Paired tumor and matched normal BAM files were analyzed with Mutect2 against the human reference genome hg38 (GRCh38). To reduce false-positive calls, a panel of normals and the gnomAD allele frequency resource were incorporated during variant calling. Raw somatic variants identified by Mutect2 were filtered using GATK FilterMutectCalls (version 4.6.2.0). Filtered variants were subsequently normalized and decomposed into primitive alleles using BCFtools (version 1.23.1) (44), and only variants annotated with a PASS filter status were retained for downstream analyses. Functional annotation was performed using GATK Funcotator (version 4.6.2.0), and variants classified as silent, intronic, or intergenic were excluded. Additional filtering retained variants with a variant allele frequency (VAF) of at least 5% and a sequencing depth of at least 30. Variants containing reference or alternate alleles longer than 20 bp were also excluded. The remaining non-synonymous somatic variants were used for subsequent analyses.

Copy number alterations were analyzed using PureCN (version 2.8.1) (45) following the recommended workflow for targeted sequencing data. Briefly, target intervals were optimized with mappability correction, and a normal reference database was constructed from matched normal samples. Coverage profiles and both germline and somatic variants identified by Mutect2 (GATK version 4.6.2.0) were integrated to estimate allele-specific copy number states, tumor purity, and ploidy. Gene-level copy number alterations used for downstream analyses were derived from the final segmentation results.

### Statistical analysis

Statistical analyses were performed using the Mann–Whitney U test, Welch’s t-test, the chi-square test, Kaplan–Meier survival analysis, and the Cox proportional hazards model in R (version 4.4.1; R Foundation for Statistical Computing) and Microsoft Excel (Microsoft).

---

**Figure S1.**
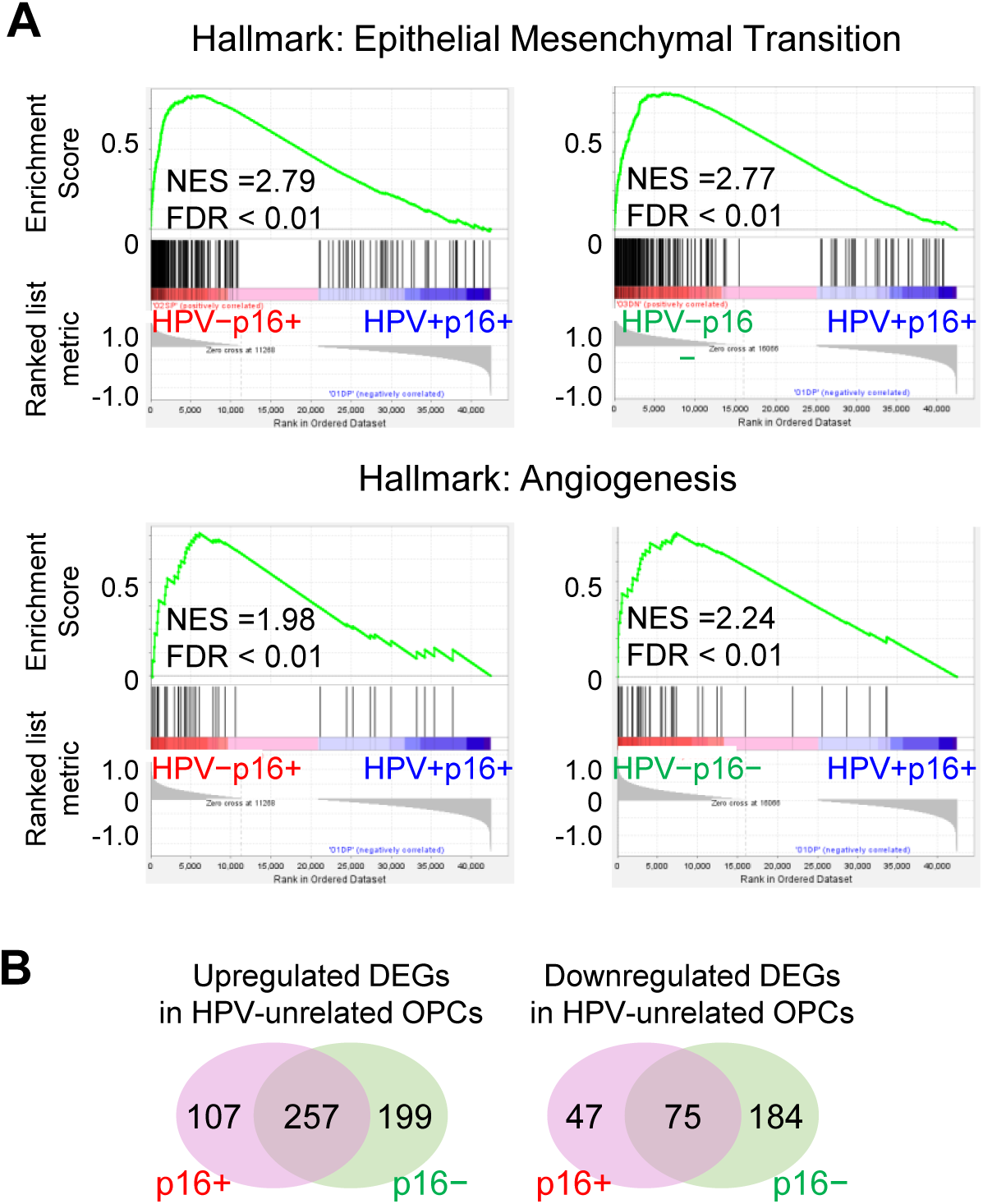
Gene expression differences between HPV-related and HPV-unrelated OPCs in our cohort. (A) GSEA plots showing gene sets upregulated in HPV-unrelated cancers compared with HPV-related cancers, irrespective of p16 status. (B) Venn diagrams showing the numbers of genes significantly upregulated or downregulated in p16^+^ and p16^-^ HPV-related cancers.

**Figure S2.**
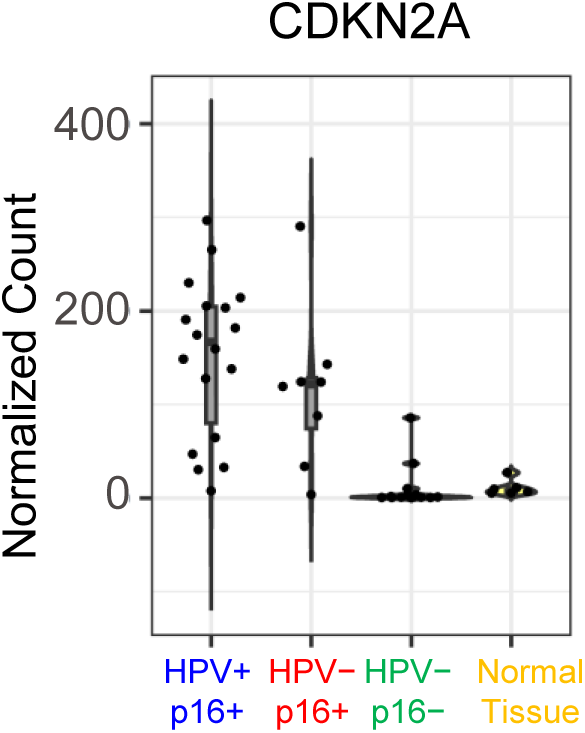
Normalized CDKN2A transcript counts from RNA sequencing according to HPV/p16 status in our cohort.

**Figure S3.**
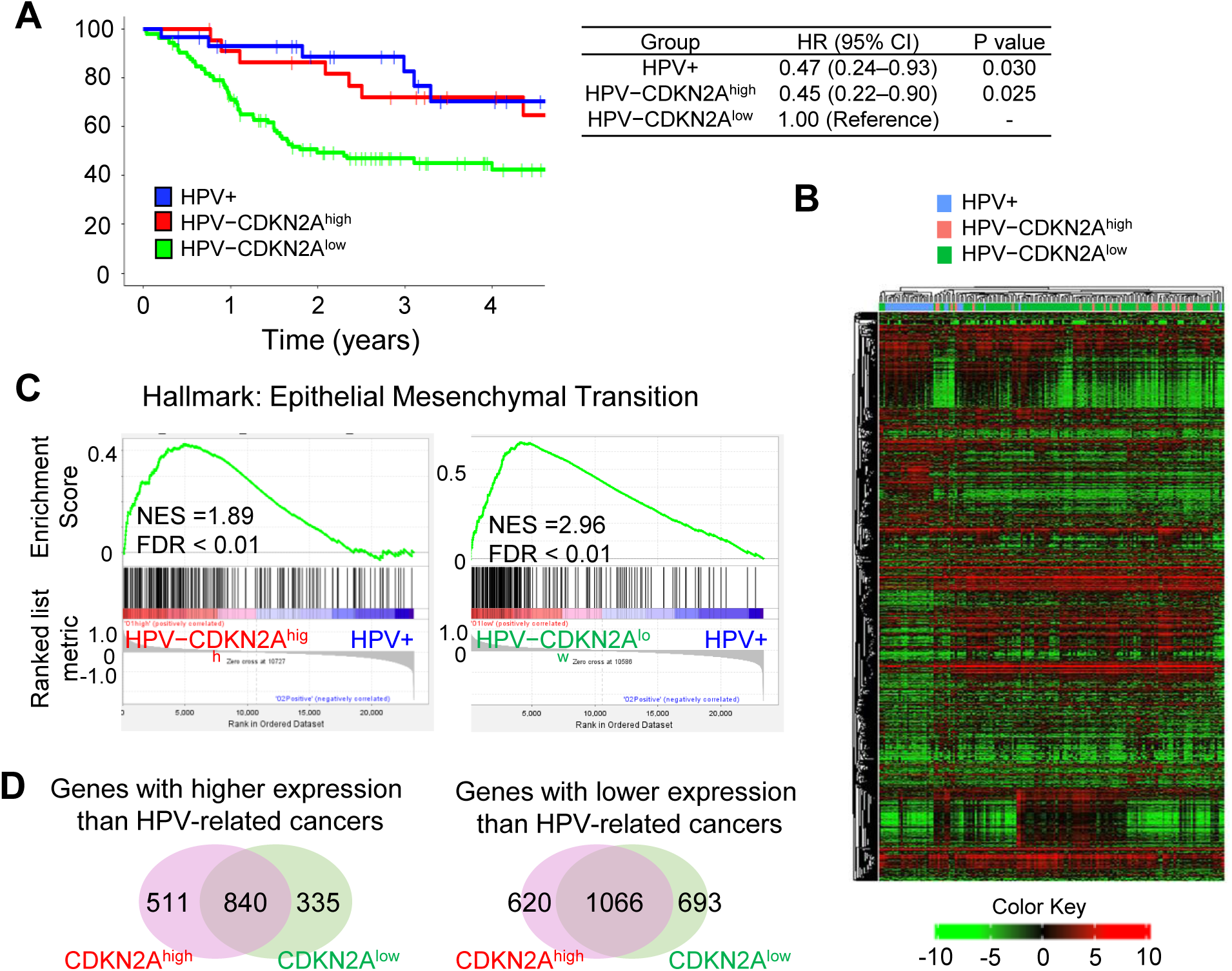
Transcriptome analysis in the TCGA dataset. (A) Kaplan–Meier curves for overall survival in TCGA cases. P values were calculated using the log-rank test. (B) Hierarchical clustering based on transcriptome data. (C) GSEA plots for the EMT gene set, upregulated in HPV-unrelated versus HPV-related cancers regardless of CDKN2A expression status. (D) Venn diagrams showing the numbers of genes significantly upregulated or downregulated in CDKN2A^high^ and CDKN2A^low^ HPV-related cancers.

**Figure S4.**
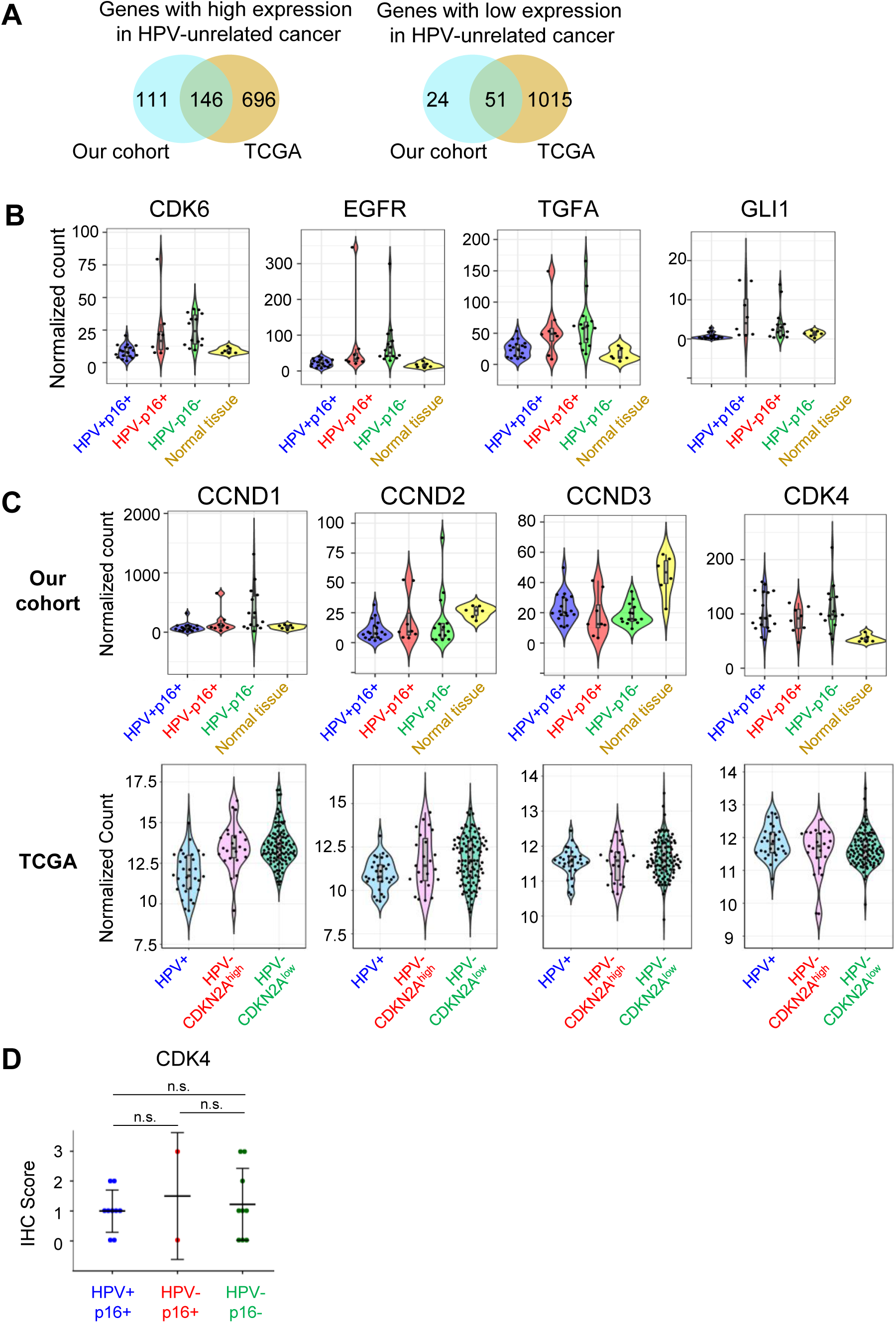
Differential gene expression between HPV-related and HPV-unrelated cancers. (A) Venn diagrams showing the overlap of genes significantly upregulated or downregulated in HPV-unrelated versus HPV-related cancers. (B) Violin plots of RNA-seq data from our cohort showing representative genes that are upregulated in HPV-unrelated OPC compared with HPV-related OPC and are annotated in the “Pathways in cancer” pathway. (C) Violin plots of RNA-seq data from our cohort and the TCGA dataset. (D) IHC scores for CDK4 staining. n.s., not significant.

**Figure S5.**
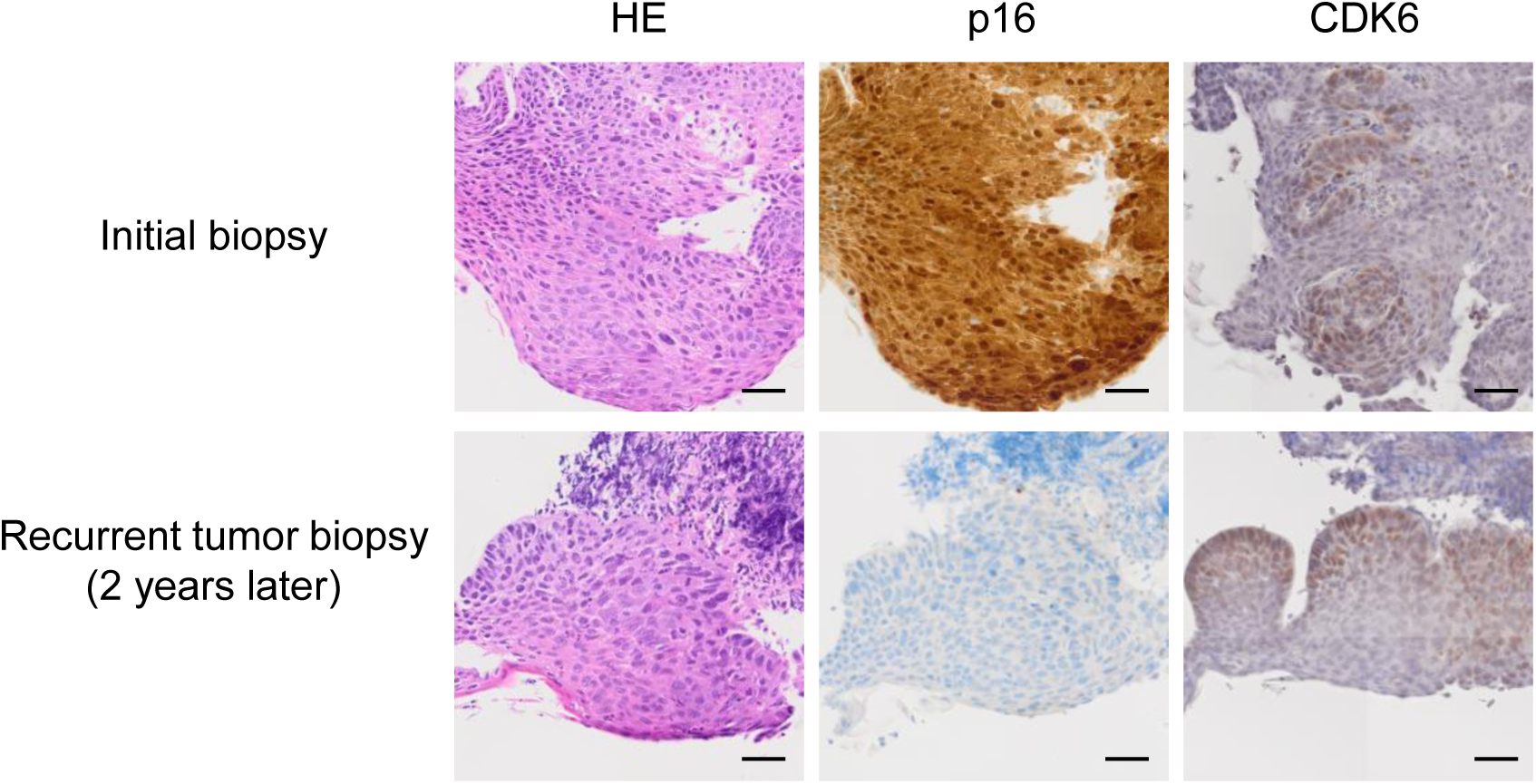
HE and IHC images of a case in which p16 staining became negative during follow-up. Scale bars, 50 μm.

**Figure S6.**
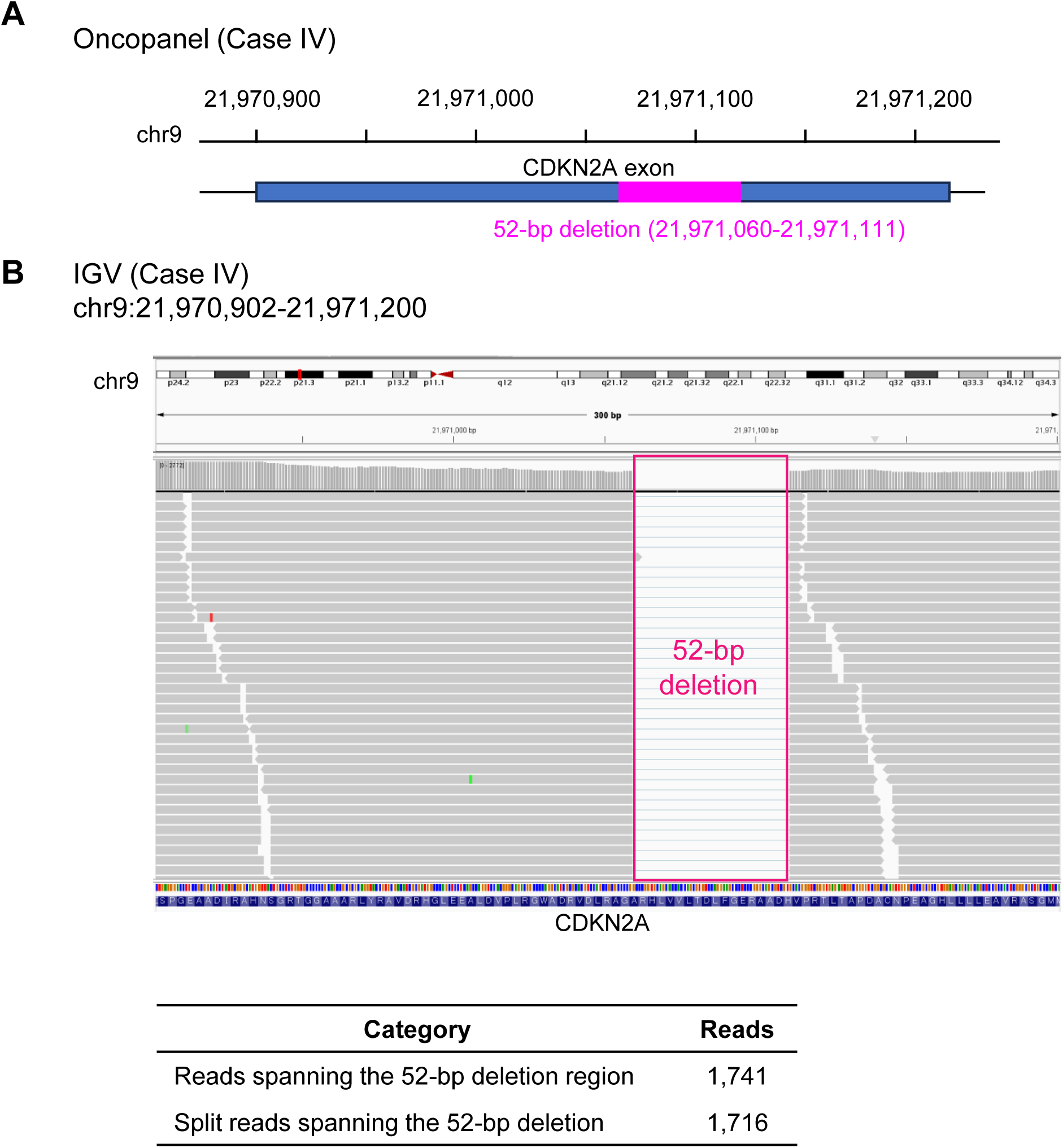
IGV view of transcriptome sequencing read alignments spanning the CDKN2A locus. (A) Schematic illustration of the CDKN2A alteration inferred from oncopanel sequencing in Case IV shown in Fig. 6B. (B) IGV view of RNA sequencing read alignments flanking the CDKN2A deletion in the same case.

**Figure S7.**
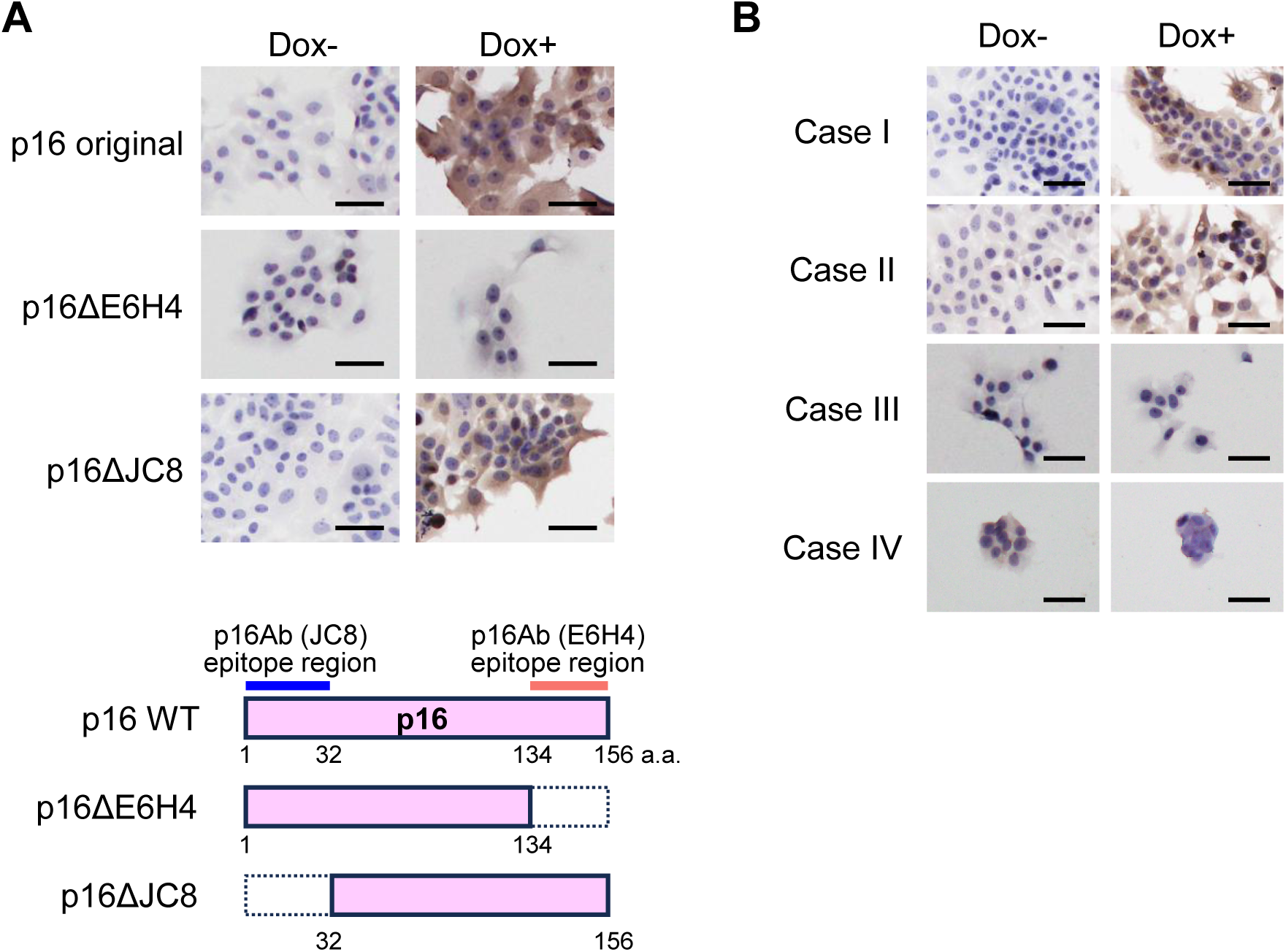
Immunostaining with an anti-p16 antibody (clone E6H4) in CDKN2A-deficient FaDu cells. (A) Wild-type p16, a C-terminal deletion mutant of p16, and a p16 construct retaining the E6H4 epitope were expressed under doxycycline control and analyzed by immunostaining. (B) Mutant p16 proteins identified in our cohort were expressed under doxycycline control and analyzed by immunostaining. The mutant labels are the same as those used in Figure 6B.

## Supplementary Tables

Table S1. Clinicopathological characteristics of OPCs in our cohort

Table S2. Hallmark gene sets upreulated or downregulated in HPV-unrelated p16-positive OPCs compared with HPV-related OPCs

Table S3. Hallmark gene sets upreulated or downregulated in HPV-unrelated p16-negative OPCs compared with HPV-related OPCs

Table S4. DEGs commonly upregulated in HPV-unrelated p16-positive and p16-negative OPCs compared with HPV-related OPCs

Table S5. DEGs commonly downregulated in HPV-unrelated p16-positive and p16-negative OPCs compared with HPV-related OPCs

Table S6. KEGG pathway enrichment analysis of DEGs commonly upregulated or downregulated in HPV-unrelated OPC

Table S7. Gene Ontology enrichment analysis of DEGs commonly upregulated or downregulated in HPV-unrelated OPC

Table S8. Hallmark gene sets upreulated or downregulated in HPV-unrelated CDKN2A-high cancers compared with HPV-related cancers

Table S9. Hallmark gene sets upreulated or downregulated in HPV-unrelated CDKN2A-low cancers compared with HPV-related cancers

Table S10. DEGs commonly upregulated in HPV-unrelated CDKN2A-high and CDKN2A-low OPCs compared with HPV-related OPCs

Table S11. DEGs commonly downregulated in HPV-unrelated CDKN2A-high and CDKN2A-low OPCs compared with HPV-related OPCs

Table S12. KEGG pathway enrichment analysis of DEGs upregulated or downregulated in HPV-unrelated OPC identified in the TCGA dataset

Table S13. DEGs significantly upregulated or downregulated in HPV-unrelated p16-positive OPCs compared with p16-negative OPCs identified in our cohort

Table S14. DEGs significantly upregulated or downregulated in HPV-unrelated CDKN2A-high OPCs compared with CDKN2A-low OPCs identified in the TCGA dataset

Table S15. Hallmark gene sets upreulated or downregulated in HPV-unrelated p16-positive OPCs compared with p16-negative OPCs in our cohort

Table S16. Hallmark gene sets upreulated or downregulated in HPV-unrelated CDKN2A-high cancers compared with CDKN2A-low cancers in the TCGA dataset

Table S17: Genes consistently upregulated or downregulated in HPV-unrelated OPCs compared with HPV-related OPCs across our cohort and the TCGA dataset

